# Study on physicochemical properties and microbial diversity of flammulina velutipes residue during rapid fermentation

**DOI:** 10.1101/428094

**Authors:** Guangying Shi, Yuxin Wang, Pingzhi Wang, Yizhu Gu

**Affiliations:** College of Water Resources & Civil Engineering, China Agricultural University, Tsinghua East Road, Haidian District, Beijing, 100083, China

**Author notes:** Corresponding author: Yuxin Wang, Ph.D., associate professor, mainly engaging in agricultural biology environment and energy engineering. Address: Tsinghua East Road, Haidian District, Beijing 100083.

**Keywords:** Flammulina velutipes, Residue, Rapid fermentation, Microbial diversity, Physicochemical properties

## Abstract

In order to investigate the effects of different mass ratios of corn straw, super absorbent resin (SAR) and cellulose decomposing strains on fermentation of flammulina velutipes residue, the cellulose degradation rate, germination index, bacterial diversity, urease activity, cellulase activity and other indicators were evaluated comprehensively so as to determine the optimal fermentation parameters. The research results indicated that the three factors of corn straw, high water-absorbent resin (SAR), and cellulose-decomposing strains have the tendency to enhance fermentation in the process. In the orthogonal test, the treatment with the highest cellulose degradation rate was T6. By 24 days, all the treated seeds germination Indices (GIs) were higher than 80%, which indicated that they were basically harmless to crops. The cellulase activity and urease activity of each treatment showed the characteristics of first rising and then decreasing as the fermentation time prolonged. In general, the T6 (the added amount of corn straw was 10%, the amount of super absorbent resin was 0.15%, and the amount of cellulose-decomposing strains was 2%) was the suitable mass ratio of additives in the fermentation which provided a certain theoretical support for resource utilization of flammulina velutipes residue.

## 0 Introduction

The development of large-scale mushroom industry has brought a great quantity of output of mushroom residue. The accumulation of mushroom residues after harvesting has become an environmental problem that cannot be ignored. If not properly handled, it will cause serious harm to our ecological environment. An estimated 2 million tons of spent mushroom substrate (SMS) are produced yearly in China, and the economic potential of recycling is significant ^[1]^. However, most of the SMS have been burnt for energy or thrown away, which is neither environment friendly nor economic ^[2]^.

The mushroom residues is a residual material remaining after the harvest of mushrooms^[3]^. The mushroom residues, which are lignocellulosic byproducts of mushroom cultivation, are hard to degrade. They usually contain high organic matter content with low macro- and micro-nutrient concentrations ^[4]^. On average, approximately 5 kg of mushroom residues is generated for each kilogram of mushroom produced ^[5]^After mushrooming, the composition of mushroom residues changed greatly. About 1/3 of the part decomposed by hyphae was used for bacterial synthesis, 1/3 was used for respiratory consumption, and the other 1/3 was present in new form in mushroom residue. Therefore, there are a large amount of nutrients that are not utilized in the mushroom residues, which contains a large number of microorganisms, bacterial proteins, various metabolites and other nutrients^[6]^such as nitrogen and phosphorus elements^[7]^ etc.. The content of organic matter in mushroom residue is 60 times that of greenhouse soil, and the total nitrogen is 100 times ^[8]^. Meanwhile, fresh mushroom residues may have beneficial effects as an organic material for soil amendment ^[9–10]^·

Moreover, studies have shown that the fermented mushroom residue is better than the unfermented mushroom residue substrate on the effect of seedling raising ^[11]^. Further treatment of mushroom residues can be potentially used as environment-friendly vegetable substrate to promote the sustainable development of agriculture ^[12, 13]^.

However, mushroom residues contain a large number of microbial communities [^14,15]^, which also carry organic acids, phenols and pathogenic bacteria and have certain toxic effects on crop growth and development ^[16]^.Therefore, mushroom residues cannot be used as cultivation substrate to grow crops directly, and it needs to be processed through fermentation. At present, there are two main ways of affecting fermentation. One is changing variables of environment, such as proper temperature, moisture content, C/N ^[17, 18]^, etc. and the second is adding additives such as corn straw and cellulose-decomposing strains to promote the process of fermentation ^[19]^, which can reduce organic acids and cellulose content^[20]^, soil-borne diseases ^[21]^, and eliminate pathogenic bacterial, regulate pH and EC value ^[22]^. Supposing that mushroom residues can be used as a re-production ingredient, it can reduce environmental pollution, replace non-renewable peat resources, and provide new ways for sustainable development.

Studies have shown that the growth of plants and the life activities of microorganisms require water as the basis^[23]^. During the fermentation process, a plastic film is placed on the stack to keep water concentration of mushroom residue at the range of 50–60%. Nevertheless, the internal oxygen concentration in the mushroom residue will be restricted by this water retention method. Therefore, in this experiment, the plastic film is replaced by adding super absorbent resin (SAR) to maintain sufficient water during the fermenting period.

In this paper, the influence of additives on flammulina velutipes residue were investigated by adding corn straw, super absorbent resin (SAR) and cellulose decomposing strains in the process of fermentation. Through high-throughput sequencing technology, single-factor tests and orthogonal tests etc., the fiber degradation rate, bacterial diversity, enzyme activity and other indicators were comprehensively evaluated to optimize the fermentation parameters of flammulina velutipes residue, which can provide a scientific basis for the reuse of flammulina velutipes residue.

## 1 Materials and Methods

### 1.1 Materials

Flammulina velutipes residue (obtained from Beijing Hualuo Biotechnology Co., Ltd.), corn straw (Zhongnong Fortis Science Park), cellulose decomposing strains (Zhongnong Lvkang Bio-Technology Co., Ltd.) and super absorbent resin (SAR, Beijing Shenglanlin Ecological Technology Co., Ltd.) were employed in the experiments. The main nutrient contents of flammulina velutipes residue and corn straw are shown in Table 1.

**Table 1:** The nutrient content of flammulina velutipes residue (mass fraction).

| Raw material | Organic matter<br>(%) | Total carbon<br>(%) | Total nitrogen<br>(%) | Total phosphorus<br>(%) | Total potassium<br>(%) |
| --- | --- | --- | --- | --- | --- |
| Corn straw | 87.16±0.54 | 46.6 | 0.9 | 0.27 | 0.59 |
| flammulina<br>velutipes residue | 85.15±0.18 | 44.8 | 1.22 | 0.47 | 1.27 |

### 1.2 Experimental design

The experiments of fermentation are divided into two parts: single factor tests and orthogonal tests. Firstly, the three levels of corn straw, super absorbent resin (SAR) and cellulose decomposing strains are selected through single-factor test. Then, data of three factors and three levels are brought into the orthogonal table, and the orthogonal test is performed. The degradation rate, germination index, cellulase activity, and urease activity are used as assessment indicators to determine optimal degradation conditions.

**Single-factor experimental design:** Flammulina velutipes residue being used as the main fermentation material, the additive contents of corn straw (A), Super Absorbent Resin (SAR) (B) and cellulose decomposing strains (C) were shown at table 2.

**Table 2:** The additive contents of single factor experiment in the fermentation.

| treatment | Corn straw A (%) | SAR B (%) | Cellulose decomposing strains C (%) |
| --- | --- | --- | --- |
| 1 | 0 | 0 | 0 |
| 2 | 10 | 0.05 | 2 |
| 3 | 20 | 0.10 | 4 |
| 4 | 30 | 0.15 | 6 |
| 5 | 40 | 0.25 | 8 |

**Orthogonal experimental design**: According to the results acquired by the single factor experiment, the orthogonal table L_9_ (3^3^) is adopted and which is shown in Table 3.

**Table 3:** Orthogonal table of *L_9_* (3^3^).

| Test treatment | A (%) | B (%) | C (%) |
| --- | --- | --- | --- |
| T1 | 3 | 1 | 1 |
| T2 | 2 | 1 | 2 |
| T3 | 1 | 1 | 3 |
| T4 | 3 | 2 | 3 |
| T5 | 2 | 2 | 2 |
| T6 | 1 | 2 | 1 |
| T7 | 3 | 3 | 2 |
| T8 | 2 | 3 | 3 |
| T9 | 1 | 3 | 1 |

### 1.3 Test Measurement Items and Methods

The main parameters need to measure are pH/EC value, germination index, cellulose degradation rate (Van Soest), cellulase activity (3, 5 dinitro Salicylic acid method), urease activity (sodium phenolate-sodium hypochlorite colorimetric method) and bacterial diversity (16sRNA-High-throughput sequencing).

## 2 Analysis and discussion on single sactor experiment

### 2.1 Comparison of cellulose degradation rates

Taking cellulose degradation rate as the assessment index, single-factor test was conducted (two factors fixed, and a single variable can change) according to Table 2. The test results are shown in Fig. 1 and Fig. 2.

**Fig. 1.**
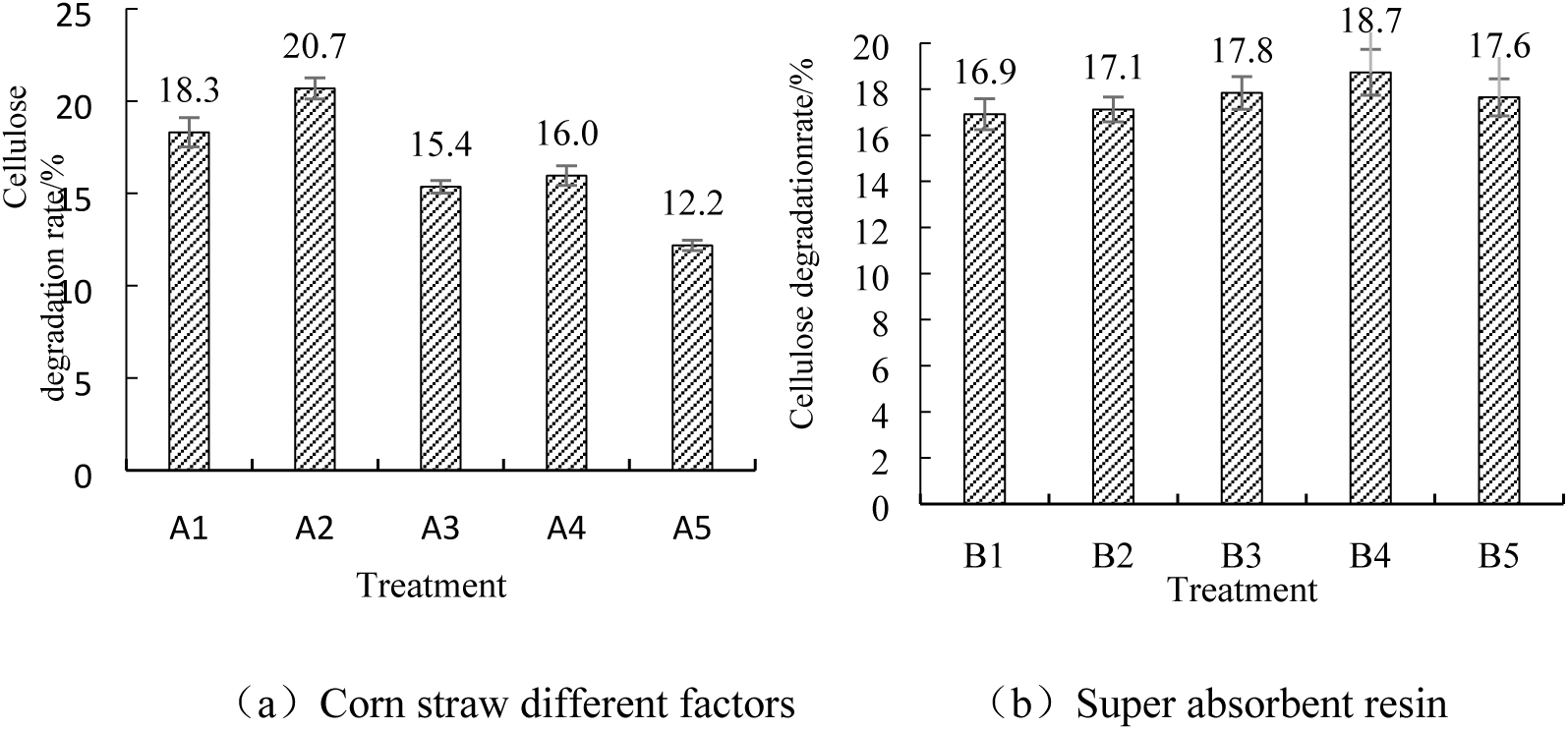
The influence of on cellulose degradation rate.

**Fig. 2.**
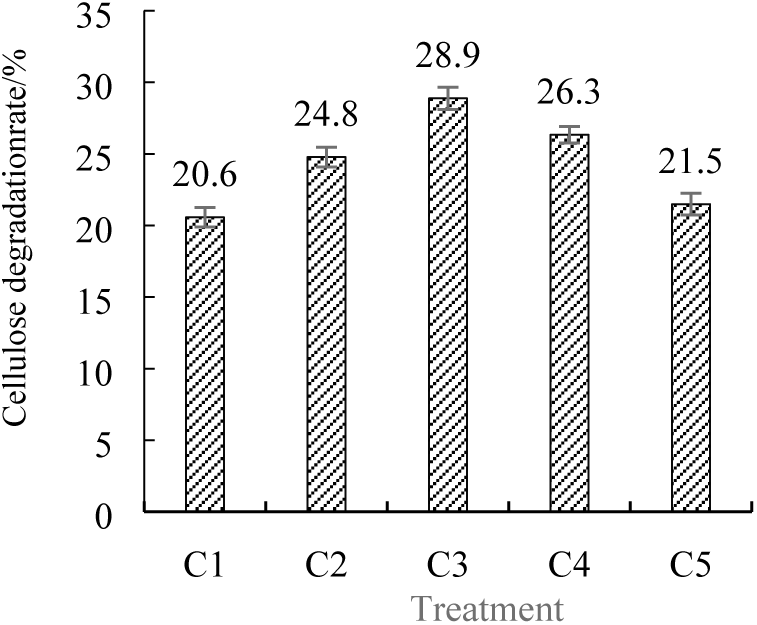
The influence of Cellulolytic bacteria on cellulose degradation rate.

Fig. 1a shows the influence of corn straw on the cellulose degradation rate. It can be seen from the figure that the degradation rate of cellulose appears a tendency of rising first and then decreasing with the increase of corn straw content. When the mass ratio of corn straw is 10%, the degradation rate of the cellulose reaches the highest.

Fig. 1b shows the influence of super absorbent resin (SAR) on the cellulose degradation rate.

It can be seen from the figure that the cellulose degradation rate shows the trend of rising first and then decreasing with the increase of the mass ratio of SAR. When the mass ratio of SAR is 0.15%, the degradation rate of flammulina velutipes residue is 18.73% at the highest. Super absorbent resin (SAR) possess a strong water retention property, it can prevent the loss of moisture during the fermenting period. However, as the increasing of SAR, it will inhibit the life activities of microorganisms and reduce the degradation rate of flammulina velutipes residue.

Fig. 2 shows the influence of cellulose-decomposing strains on the cellulose degradation rate. It can be seen from the figure that the cellulose degradation rate shows the trend of rising first and then decreasing with the increase of cellulose-decomposing strains. When the mass ratio of added cellulose-decomposing strains rising to 4%, the degradation rate of cellulose in flammulina velutipes residue reached at 28.88% being the highest.

### 2.2 Influence of additives on bacterial diversity

In order to explore the diversity of microorganisms in the experiments, 4 test groups (n=3 in each group) were arranged separately. They are GZ_CS, GZ_T1, GZ_T2 and GZ_T3, where CS is the control group. Fig. 2 The influence of Cellulolytic bacteria on cellulose degradation rate. In each group, plastic pots (height, 16 cm; diameter, 12.5 cm) were respectively filled with fermenting flammulina velutipes residue and additive contents (corn straw, super absorbent resin (SAR) and cellulose decomposing strains) were shown in the table 4. In the group GZ_CS, only flammulina velutipes residue is included. GZ_T1 is the group of adding SAR, GZ_T2 is the group of adding cellulose decomposing strains, and GZ_T3 is the group of adding corn straw.

**Table 4:**
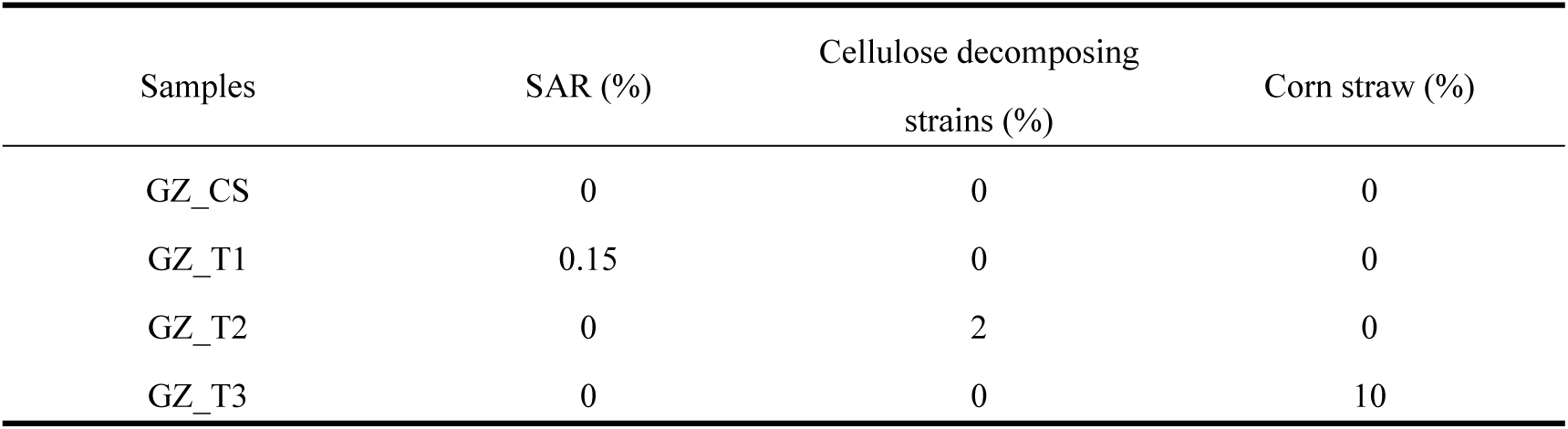
Mass ratios of additives in the experimental groups.

From Table 5, it can be seen that the Shannon index of groups GZ_T1, GZ_T2 and GZ_T3 are higher than that of GZ_CS, which indicated that the additives of SAR, corn straw, and cellulose-degrading strains can promote the growth of bacteria effectively. Microorganisms are abundant in every group, and the order of microbial population abundance is GZ_T2>GZ_T3>GZ_T1.

**Table 5:** Bacterial diversity indices.

| Samples | Similarity level=0.97 |  |  |  |  |  |
| --- | --- | --- | --- | --- | --- | --- |
|  | Sobs | Shannon | Simpson | Ace | Chao | Coverage |
| GZ_CS | 107 | 2.7736 | 0.1213 | 113.2533 | 109.7692 | 0.9998 |
| GZ_T1 | 238 | 3.2976 | 0.0777 | 250.7006 | 249.0400 | 0.9994 |
| GZ_T2 | 260 | 3.6431 | 0.0549 | 286.2608 | 291.7143 | 0.9988 |
| GZ_T3 | 250 | 3.4641 | 0.0742 | 269.2111 | 272.5455 | 0.9988 |

The community richness of bacteria is characterized by Sobs index, Chao index and Ace index. Bacterial community diversity is represented by Shannon Index and Simpson Index. The larger Shannon index indicates that the diversity of the community is more abundant in the corresponding group. The larger the Simpson index, the lower the diversity of the community^[15, 16]^ in the corresponding group.

Fig. 3 shows the species composition at the level of phylum and genus of bacterial colonies in each group after 24 days’ treatment. Fig. 3a shows that *Actinobacteria* and *Bacteroidetes* in the groups of GZ_T1, GZ_T2, and GZ_T3 are richer than that of GZ_CS. The proportions of *Actinobacteria* and *Bacteroidetes* in the group GZ_T1 are higher, which reached at 60.12% and 16.90%, respectively. The proportions of *Actinobacteria* and *Bacteroidetes* in the group GZ_T3 are 54.99% and 12.40%, respectively. It is can be seen that SAR has an enhancing effect on *Actinobacteria* and *Bacteroidetes* in the fermentation materials.

**Fig. 3.**
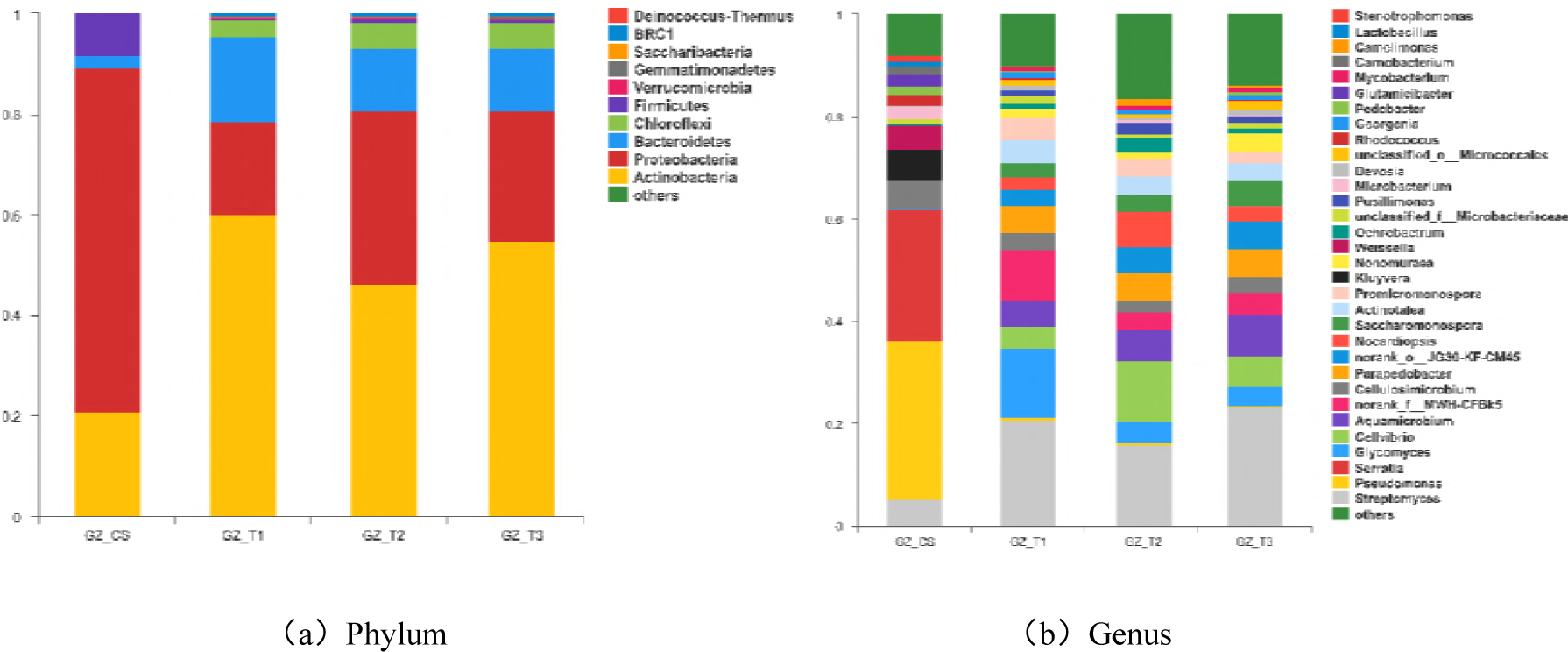
Bacterial analysis of mushroom residue after treatment under different conditions.

Fig. 3b shows that the proportions of *Streptomyces* and *Glycomyces* in the groups of GZ_T1, GZ_T2, and GZ_T3 are higher than that of the group GZ_CS. *Glycomyces* in the GZ_T1 group has the largest proportion at 13.45%. The GZ_T3 group has the highest proportion of *Streptomyces*, which is 23.37%. The proportions of *Pseudomonas* and *Serratia* in the group GZ_CS is significantly higher than that of the other three groups, which are 31.03% and 25.76%, respectively. It can be seen that after further fermentation, the proportions of *Streptomyces* and *Glycomyces* in the three groups containing superabsorbent resin, cellulolytic strains and corn straw appear some certain increases, while the proportions of *Pseudomonas* and *Serratia* decrease significantly.

After adding the three different additives in the fermentation, the species of original bacteria in the flammulina velutipes residue showed significant enhancing phenomenon. The SAR can lock the moisture in material during the decomposing process to make abundance of the microorganisms higher and keep for a relative long period. Cellulose-decomposing strains is an additive material rich of microbial population. Smashed corn straw with low density is bulky and fluffy. When it is added to the flammulina velutipes residue, the porosity of materials can be adjusted well, which can enhance the efficiency of microbial aerobic fermentation.

## 3 Orthogonal test results analysis

### 3.1 The Effect of different treatment on cellulose degradation rate

Through the single-factor test results, the three levels of corn straw (A), SAR (B), and cellulose decomposing strains (C) were arranged in Table 6.

**Table 6:** Level of orthogonal experiment.

| Level | A/% | B/% | C/% |
| --- | --- | --- | --- |
| 1 | 10 | 0.05 | 2 |
| 2 | 20 | 0.15 | 4 |
| 3 | 30 | 0.25 | 6 |

The levels in table 6 were substituted into the orthogonal table L_9_(3^3^) and the test results are shown in Table 7.

**Table 7:** Orthogonal test result of additives on flammulina velutipes residue fermentation.

| Test treatment | A/% | B/% | C/% | Cellulose degradation rate/% |
| --- | --- | --- | --- | --- |
| T1 | 30 | 0.05 | 2 | 22.70 |
| T2 | 20 | 0.05 | 4 | 22.67 |
| T3 | 10 | 0.05 | 6 | 21.01 |
| T4 | 30 | 0.15 | 6 | 22.98 |
| T5 | 20 | 0.15 | 4 | 22.59 |
| T6 | 10 | 0.15 | 2 | 25.04 |
| T7 | 30 | 0.25 | 4 | 22.23 |
| T8 | 20 | 0.25 | 6 | 21.76 |
| T9 | 10 | 0.25 | 2 | 20.99 |
| $K_1$ | 67.91 | 66.39 | 68.73 | |
| $K_2$ | 67.03 | 70.62 | 67.50 | |
| $K_3$ | 67.05 | 64.99 | 65.76 | |
| $k_1$ | 22.64 | 22.13 | 22.91 | |
| $k_2$ | 22.34 | 23.54 | 22.50 | |
| $k_3$ | 22.35 | 21.66 | 21.92 | |
| R | 0.3 | 1.88 | 0.99 |  |

K_i_ (column j) in the table above: the sum of the data in the jth level; k_i_ (column j): the comprehensive average of the jth level data; R: The difference of ki in the jth column (i=1, … 3.j=1,…3).

In addition, the results of the orthogonal test need to be analyzed by the F-test to further verify the differences among the three factors (Table 8). The order of three influence factors, from big to small is B>C>A.

**Table 8:** The ANOVA analysis of cellulose degradation rate.

| Source of variation | Sum of square(SS) | Degree of freedom(df) | Mean square(MS) | F | $F_{0.05}$ |
| --- | --- | --- | --- | --- | --- |
| A | 0.75 | 2 | 0.37 | 0.31 |  |
| B | 3.76 | 2 | 1.88 | 1.56 | 19.37 |
| C | 0.77 | 2 | 0.39 | 0.32 |  |
| Error | 2.42 | 2 | 1.21 |  |  |
| Total variation | 7.69 | 8 |  |  |  |

The analysis results showed that the three factors had no significant difference under the level *α* =0.05. Therefore, the suitable treatment can be selected from table 7, that is T6. In other words, when the additive amount of corn straw was 10% (A1), the amount of high water-absorbing resin (SBR) was 0.15% (B2), and the mass ratio of cellulose decomposing strains was 2% (C1), the degradation rate of cellulose reached at the highest, which was 25.04%.

**Fig. 4.**
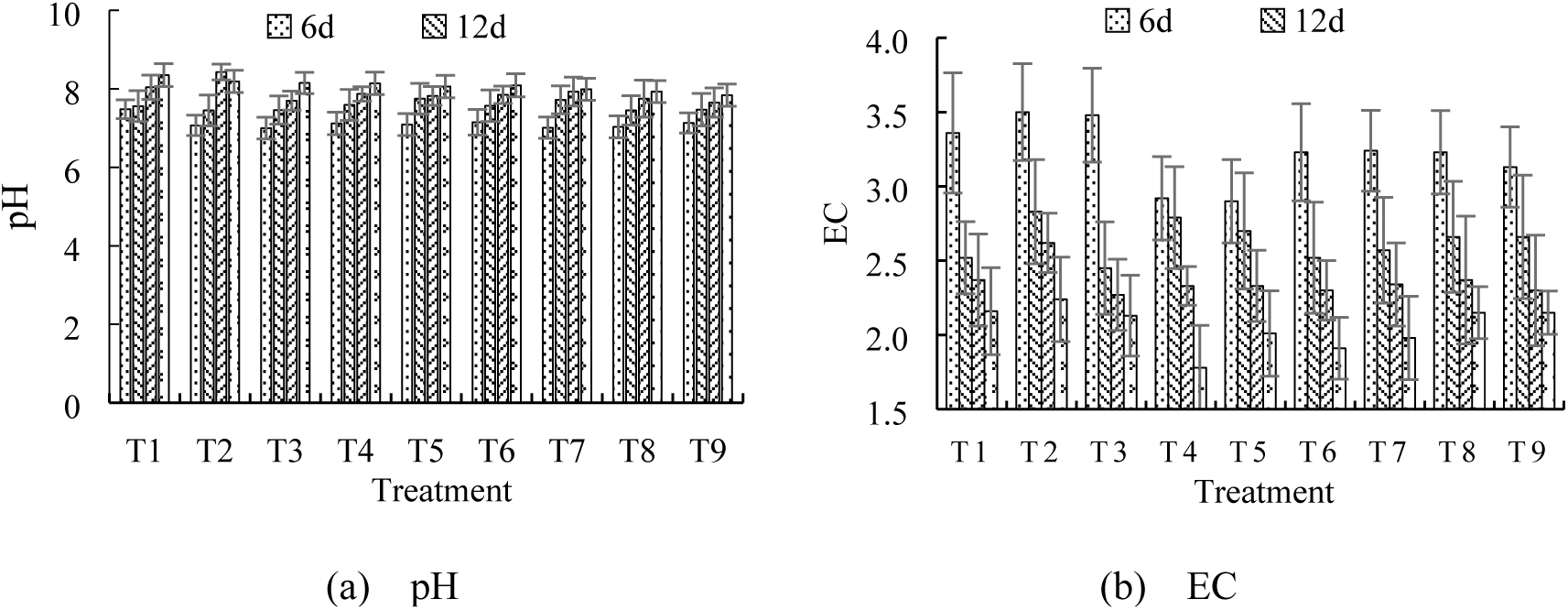
Effects of different treatments on pH and EC values of mushroom residue.

### 3.2 Effect of Different Treatment on pH, EC and GI

Fig. 4 shows the effect of each treatment on the value pH and Electric Conductivity (EC). It can be seen that during the fermentation period of 24 days, the value of pH in each treatment was gradually rising (Fig. 4a), and the EC value was decreasing sharply with the increasing of fermentation time (Fig. 4b).

Germination index (GI), as an assessment parameter of biology, it can objectively describe the toxicity of substrate to plants^[24]^. Studies have shown that when the value of GI higher than 50%, the substrate is essentially no inhibitory effect on the plant^[25]^, and when GI higher than 80%, the fermented material is completely non-toxic to the plant^[26]^.

**Fig. 5.**
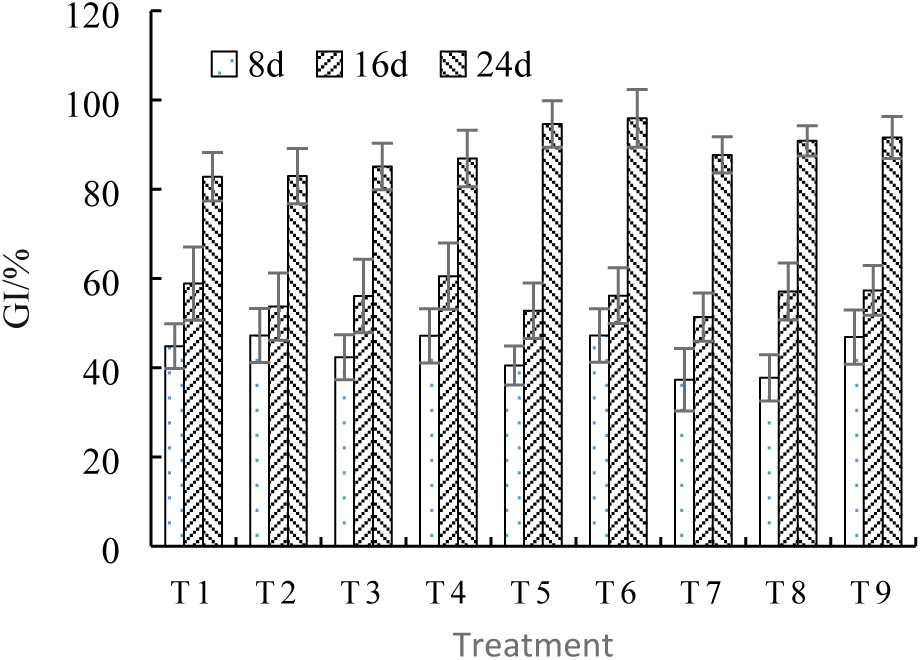
Effects of Different Treatments on GI of Cucumber.

It can be seen from Fig. 5 that at the early stages of fermentation, mushroom residues have inhibitory or even toxic effects on plant seeds, and the GI value is relatively lower. It may be that the flammulina velutipes residue can produce some toxic substances during initial fermentation, such as aldehydes, phenols and organic acids, etc.^[27]^. With the increase of fermenting days, the germination index of substrate increased gradually. On the 24th day of fermentation, the GI value rising at 80%, it indicated that long-term aerobic decomposition can improve the chemical properties of substrates.

### 3.3 Effects of different treatments on urease and cellulase activities

All biochemical changes in the process of fermentation are carrying on with the participation of various enzymes. Figure 6a is the dynamic changes of urease activity under different treatments. On the eighth day the urease activity of each treatment was lower, and it was higher at the sixteenth days. On the twenty-fourth day of fermentation, the urease activity was greatly decreased. Taking T7 as an example, the urease activity reached 262.66 U/g at sixteenth days after fermentation, and at the twenty-fourth day of fermentation, it decreased to about 100 U/g.

Cellulase is a general term for a group of enzymes which can degrade cellulose to glucose. It is not a monomer enzyme, but a synergistic multicomponent enzyme system. As biocatalysts in the decomposition of cellulose, cellulase can break down cellulose into oligosaccharides or monosaccharides. Fig. 6b shows the effect of different treatments on the cellulase activity. The cellulase activity had the tendency of rising first and then decreasing during the fermentation period. Taking T6 as an example, the cellulase activity was about 65 U/g on the eighth day, and at the sixteenth day of fermentation the cellulase activity reached 95 U/g. At the twenty-fourth day of fermentation, it decreased to about 30 U/g.

**Fig. 6.**
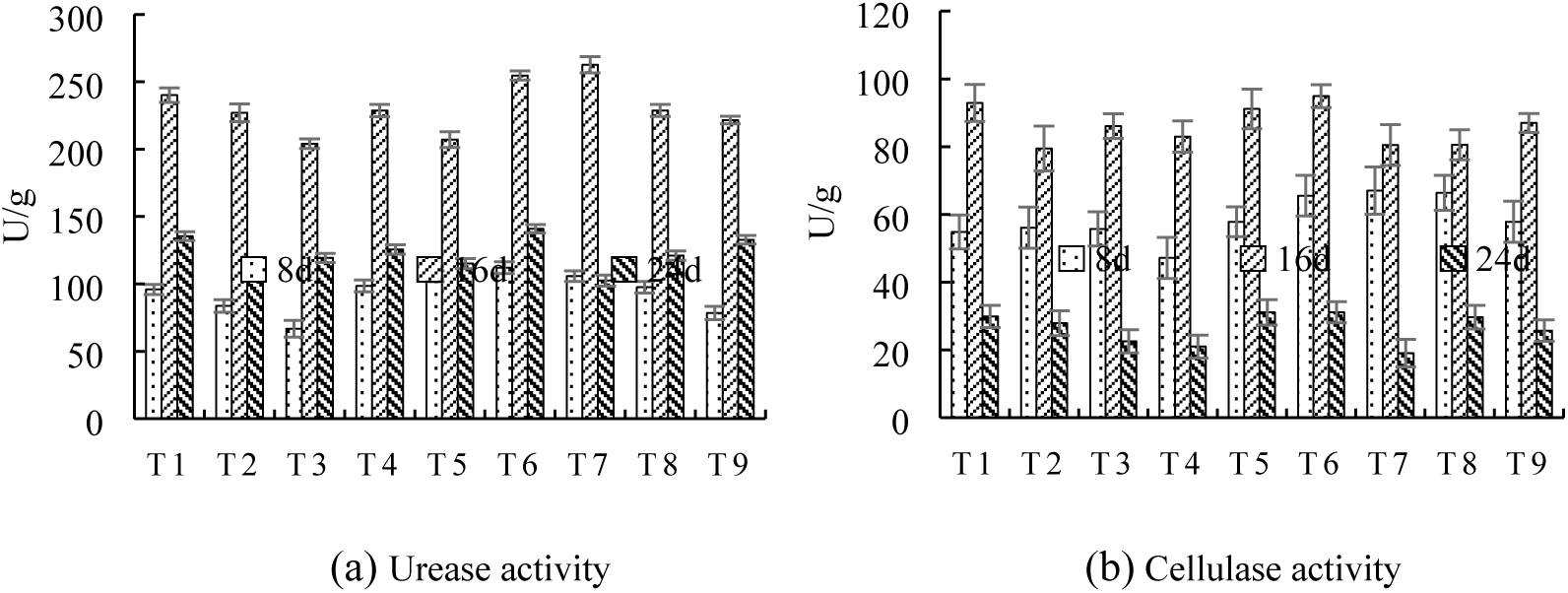
Effects of different treatments on enzyme activity.

## 4 Conclusion

Technical progress of high-throughput sequencing has allowed researchers to reveal the microbial diversity at an unprecedented level compared with the traditional cultural-based and PCR-DGGE way. Using high-throughput sequencing technology, it is more intuitively and detailedly understand the effects of different additives on community structure and species composition in the flammulina velutipes residue. The results of high-throughput sequencing indicate that Actinobacteria and Bacteroidetes in the groups of GZ_T1, GZ_T2, and GZ_T3 are richer than that of the control group GZ_CS. The proportions of *Streptomyces* and *Glycomyces* in the groups of GZ_T1, GZ_T2, and GZ_T3 are higher than that of the GZ_CS group. The proportions of *Pseudomonas* and *Serratia* in the group GZ_CS are significantly higher than that of the other three groups. In general, the additives of corn straw, high water-absorbent resin (SAR) and cellulose-degrading strains all have a certain role in promoting the fermentation of flammulina velutipes residue.

The orthogonal test showed that the treatment with the highest cellulose degradation rate was T6 group. It means the optimum mass ratios of SAR, cellulose decomposing strains and corn straw in the fermentation of flammulina velutipes residue w 0.15%, 2% and 10%, respectively. on the 24th day of fermentation, the GI value rising at 80% in each treatment, it indicated that long-term aerobic decomposition can improve the chemical properties of substrates.

**Data availability statement** The [Raw sequence reads] data used to support the findings of this study have been deposited in the [NCBI] repository (Accession: Accession: PRJNA474591)

## References

[1] Qiao, J.J., Zhang, Y.F., Sun, L.F., Liu, W.W., Zhu, H.J., Zhang, Z.J. Production of spent mushroom substrate hydrolysates useful for cultivation of Lactococcus lactis by dilute sulfuric acid, cellulase and xylanase treatment. Bioresour Technol, 2011,102, 8046–8051.

[2] Zhu, H.J., Liu, J.H., Sun, L.F., Hu, Z.F., Qiao, J.J. Combined alkali and acid pretreatment of spent mushroom substrate for reducing sugar and biofertilizer production. Bioresour. Technol, 2013,136, 257–266.

[3] Phan, C.-W., Sabaratnam, V.. Potential uses of spent mushroom substrate and its associated lignocellulosic enzymes. Appl. Microbiol. Biotechnol, 2012,96, 863–873

[4] Medina, E., Paredes, C., Bustamante, M.A., Moral, R., Moreno-Caselles, J..Relationships between soil physico-chemical, chemical and biological prop-erties in a soil amended with spent mushroom substrate. Geoderma, 2012,152–161.

[5] Lau, K.L., Tsang, Y.Y., Chiu, S.W. Use of spent mushroom compost to bioremediate PAH-contaminated samples. Chemosphere, 2003,52, 1539–1546.

[6] Buswell J.A. Potential of spent mushroom substrate for bioremediation purposes[J]. Compost Science and Utilization, 1994,2(3):31–36.

[7] Lou Zimo, Sun Yue, Zhou Xiaoxin, et al. Composition variability of spent mushroom substrates during continuous cultivation, composting process and their effects on mineral nitrogen transformation in soil[J]. Geoderma, 2017,307:30–37.

[8] Polat E, Uzun H I, Top□uoglu B, et al. Effects of spent mushroom compost on quality and productivity of cucumber (Cucumis sativus L.) grown in greenhouses [ J]. African Journal of Biotechnology,2010,8 (2): 176 — 180

[9] Hiroko Nakatsuka, Masato Oda, Yukimi Hayashi, Kenji Tamura. Effects of fresh spent mushroom substrate of Pleurotus ostreatus on soil micromorphology in Brazil[J]. Geoderma,2016,269.

[10] Oda, M., Tamura, K., Nakatsuka, H., Nakata, M., Hayashi, Y. Application of high carbon:nitrogen material enhanced the formation of the soil a horizon and nitrogen fixation in a tropical agricultural field. Agric. Sci. 5, 2014, 1172–1181

[11] Hong Chunlai, Wang Weiping, Chen Xiaotong, et al. Application of mushroom slag composite matrix in cucumber seedlings [J]. Zhejiang Agricultural Science, 2010 (6):1203–1206.

[12] Medina E., Paredes C., Perez-Murcia M.D., et al. Spent mushroom substrates as component of growing media for germination and growth of horticultural plants[J]. Bioresource Technology, 2009,100(18):4227–4232.

[13] Vellinga E.C. The mushroom monitoring project of the future: A plea for much more ecological research.[J]. Coolia, 2012,55(1):7–12.

[14] McGee Conor F. Microbial ecology of the Agaricus bisporus mushroom cropping process[J]. Applied Microbiology and Biotechnology, 2018,102(3):1075–1083.

[15] Xiao Zheng, Lin Manhong, Fan Jinlin, et al. Anaerobic digestion of spent mushroom substrate under thermophilic conditions: Performance and microbial community analysis[J]. Applied Microbiology and Biotechnology, 2018,102(1):499–507.

[16] Paula Fabiana S., Tatti Enrico, Abram Florence, et al. Stabilisation of spent mushroom substrate for application as a plant growth-promoting organic amendment[J]. Journal of Environmental Management, 2017,196:476–486.

[17] Yan Zhiying, Song Zilin, Li Dong, et al. The effects of initial substrate concentration, C/N ratio, and temperature on solid-state anaerobic digestion from composting rice straw[J]. Bioresource Technology, 2015,177:266–273.

[18] Estevez Maria M., Linjordet Roar, Morken John. Effects of steam explosion and co-digestion in the methane production from Salix by mesophilic batch assays[J]. Bioresource Technology, 2012,104:749–756.

[19] Niu Mingfen, Pang Xiaoping, Chen Shiruo. The study of influencing factors to corn straw mixed with pig effluent anaerobic fermentation[M]//Dan Y. Procedia Environmental Sciences. 2011: 54–60.

[20] Vuorinen A. H., Saharinen M. H. Evolution of microbiological and chemical parameters during manure and straw co-composting in a drum composting system[J]. Agriculture Ecosystems & Environment, 1997,66(1):19–29.

[21] Aziz R. A. T. A., Al-Barakah F. B. N. Composting technology and the impact of compost on soil biochemical properties[J]. Arab Gulf Journal of Scientific Research, 2005,23(2):80–91.

[22] Raviv Michael. The use of compost in growing media as suppressive agent against soil-borne diseases[J]. Acta Hort, 2008(779):39–50.

[23] Arbona V., Iglesias D.J., Jacas J., et al. Hydrogel substrate amendment alleviates drought effects on young citrus plants[J]. Plant and Soil, 2005,270(1–2):73–82.

[24] Pampuro N., Santoro E., Cavallo E. Evaluation of maturity and fertilizer capacity of compost derived from swine solid fraction.[M],2010:28.

[25] Zucconi F., Monaco A., Debertoldi M. Biological evaluation of compost maturity[J]. Biocycle, 1981,22(4):27–29.

[26] Tiquia S.M., Tam NFY, Hodgkiss I.J. Effects of composting on phytotoxicity of spent pig-manure sawdust litter[J]. Environmental Pollution, 1996,93(3):249–256.

[27] Ofosu-Budu G.K., Hogarh J.N., Fobil J.N., et al. Harmonizing procedures for the evaluation of compost maturity in two compost types in Ghana[J]. Resources Conservation and Recycling, 2010,54(3):205–209.

